# Criticality as a measure of developing proteinopathy in engineered human neural networks

**DOI:** 10.1101/2020.05.03.074666

**Authors:** Vibeke D. Valderhaug, Kristine Heiney, Ola Huse Ramstad, Geir Bråthen, Wei-Li Kuan, Stefano Nichele, Axel Sandvig, Ioanna Sandvig

## Abstract

A patterned spread of proteinopathy represents a common characteristic of many neurodegenerative diseases. In Parkinson’s disease (PD), misfolded forms of alpha-synuclein proteins aggregate and accumulate in hallmark pathological inclusions termed Lewy bodies and Lewy neurites, which seems to affect selectively vulnerable neuronal populations and propagate within interconnected neuronal networks. Research findings suggest that these proteinopathic inclusions are present at very early timepoints in disease development, even before strong behavioural symptoms of dysfunction arise, but that these underlying pathologies might be masked by homeostatic processes working to maintain the function of the degenerating neural circuits. This study investigates whether inducing the PD-related alpha-synuclein pathology in engineered human neural networks can be associated with changes in network function, and particularly with network criticality states. Self-organised criticality represents the critical point between resilience against perturbation and adaptational flexibility, which appears to be a functional trait in self-organising neural networks, both in vitro and in vivo. By monitoring the developing neural network activity through the use of multielectrode arrays (MEAs) for a period of three weeks following proteinopathy induction, we show that although this developing pathology is not clearly manifest in standard measurements of network function, it may be discerned by differences in network criticality states.

## Introduction

Neurodegenerative diseases, such as Parkinson’s disease (PD), Alzheimer’s disease (AD) and amyotrophic lateral sclerosis (ALS), represent a common cause of morbidity and cognitive impairments in older adults. Although characterised through complex pathologies and unknown aetiologies, some prominent commonalities, such as the presence of proteinopathy and the patterned spread of pathology through selectively vulnerable neuronal populations, cannot be ignored (1-9). Focusing on the second most common neurodegenerative disease, PD, the implicated proteinopathy mainly consists of misfolded and aggregated forms of alpha-synuclein. These intracellular alpha-synuclein inclusions are termed Lewy bodies or Lewy neurites, and can be found propagating throughout central, peripheral and autonomic parts of the nervous system, as well as in multiple organs, as the disease progresses (6, 7, 10-14). Furthermore, the neurodegenerative process characteristic of PD particularly affects and progressively depletes the dopaminergic neurons in the substantia nigra pars compacta (SNpc), a process which is thought to underlie most of the movement-related symptoms (15).

The neurodegenerative process underlying PD inevitably affects both the structural and functional connectivity of local and distal circuitry in the brain. As already noted, the movement-related impairments in PD are largely ascribed to the characteristic loss of dopaminergic neurons in the SNpc, and the degeneration of the nigrostriatal (dorsal striatal) pathway. However, the disease is much more systemic, affecting both the mesolimbic dopaminergic (ventral striatal) pathway and the mesocortical pathway, as well as several other neuronal populations throughout the brain as it progresses (16). As clusters of neurons and their interconnections degenerate, homeostatic plasticity mechanisms likely compensate to maintain stable function through regulation and rearrangement of synapses and synaptic elements in local and distal neural networks (17-27). The network disturbances likely imposed by the degeneration of neurons and their interconnections are thus counterbalanced, keeping clear symptoms of dysfunction from arising at early stages of disease development and thus patients from being diagnosed before advanced neurodegeneration is already present (28-30).

Interestingly, the high interconnectivity of the brain has been shown to inherently shape how it responds to perturbation, and the particular sites affected, as well as their level of connectedness to other brain regions, to determine how pathology can spread (31-34). This relates directly to the hallmark pathology of PD, namely the widespread alpha-synuclein inclusions, which have been suggested to propagate between anatomically highly interconnected areas in an “evolving topographical progression” (6, 7, 35). Based on this, neuroscientific research has narrowed in on two likely pathological mechanisms that could underlie the propagating pattern of neurodegeneration seen in PD (as well as in AD and ALS), namely selective neuronal vulnerability and pathological proteinopathic seeds (2, 3, 6, 33, 36-38). At this point, research efforts have uncovered several mechanisms of neuron-to-neuron transfer of pathological seeds of alpha-synuclein pre-formed fibrils (PFFs) (39-43), both in vitro and in vivo (11, 13), highlighting its contribution as a source of pathological propagation, however, the functional consequences of this interneuronal spread remain to be elucidated.

How can the functional consequences of such pathological mechanisms be studied? As mentioned, fundamental homeostatic plasticity mechanisms, which serve to maintain stable function in neural circuits in the face of perturbation, likely help mask the ongoing pathological processes underlying the progressive neurodegenerative pattern of PD (17, 18). However, although early network disturbances caused by the presence of pathological aggregates and the degeneration of neurons and their interconnections might be concealed in terms of behavioural symptoms, they may be detectable as fluctuations or deviations in some measures of the network function and activity state. As such, the universal attribute of neural network development towards the state of self-organized criticality (SoC) seems a logical point of focus and of particular interest. SoC represents the critical point between resilience against perturbation and adaptational flexibility, which appears without the need for fine-tuning of parameters through basic self-organizing processes in neural networks, both in vivo and in vitro (44-56). This dynamic state is characterized by cascades of spontaneous activity with power-law size distributions, activity which is electrophysiologically measurable and termed “neuronal avalanches”(45, 46, 49, 55, 56). As damage spreads within a neural network, it is highly conceivable that the system approaches a “damage threshold”, where restoring network function becomes increasingly difficult, and which represents a deviation from the criticality state (26, 27, 53, 57). Since SoC also appears in neural networks in vitro (58), functional network alterations resulting from induced pathology such as proteinopathy can be studied within this paradigm.

To investigate whether a developing PD-related proteinopathy can be associated with network criticality states, we have induced proteinopathy in engineered human neural networks in vitro and applied computational analysis to identify criticality in electrophysiological microelectrode array (MEA) recordings of the resulting network activity. Specifically, we measured the developing network activity prior to and for three weeks following exogenous addition of alpha-synuclein PFF seeds, and aimed to investigate how this induced PD-related pathology is reflected in several measures of network function and in the network criticality states compared to control neural networks. Our results suggest that induction of proteinopathy likely affects neural network behaviour in relation to SoC. To the best of our knowledge, this is the first study to investigate SoC in biological, human induced pluripotent stem cell (iPSC)-derived neural networks.

## Materials and methods

### Reprogramming of human iPSCs to neural progenitor cells

Human induced pluripotent stem cells (iPSCs) (ChiPSC18, Takara Bioscience) were reprogrammed using a protocol for midbrain dopaminergic neurons adapted from Kirkeby et. al 2012 (59) and 2016 (60) and Doi et al. 2014 (61) (Fig.1, Supplem.1). Briefly, the human iPSC were seeded on human recombinant laminin 111 (LN111, BioLamina) at a density of 10.000 cells/cm^2^, where they were exposed to dual-SMAD inhibition (LDN1931892 and SB43152), followed by Wnt signalling activation through the GSKβ inhibitor CHIR99021, and sonic hedgehog introduction (Shh C25ll) (day 0-9). On day 11 the cells were dissociated using accutase and reseeded on LN111 at a density of 50.000cells/cm^2^. FGF8b was added from day 9-16, at which point the reprogramming was concluded and the human iPSC-derived neurons were left for maturation.

**Fig.1.**
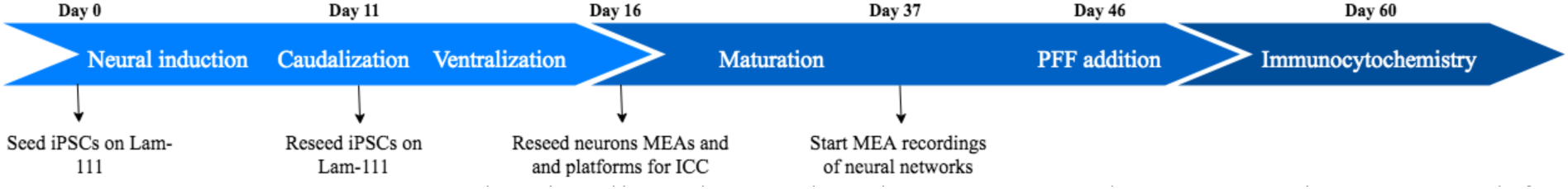
Experiment layout. The timeline shows the chemotemporal reprograming protocol for the human iPSC-derived neurons, followed by the establishment and maturation of the neural networks on multielectrode arrays (MEAs). Following 30 days of maturation, sonicated pre-formed alpha-synuclein fibrils were added to the engineered neural networks.

### Formation of alpha-synuclein pre-formed fibrils (PFFs)

Alpha-synuclein PFFs were formed following a modification of the procedure described in the protocol by Kuan et al. (62). Briefly, 1mg alpha-synuclein monomers (S-1001-1, rPeptide) was resuspended in 1mL MilliQ water, giving 1mg/mL in 20mM Tris-HCL, pH7.4, 100mM NaCl. The suspension was then centrifuged at 3600xg for 60 minutes in an Amicon Ultra 3K membrane device, which was then inverted and spun down in a tube for 1000xg for 2 minutes to transfer the concentrated sample. The concentrated solute was then resuspended to a final volume of 500ul (5mg/mL) in 10mM Tris-HCL (1.576g/L), pH 7.6, 50mM NaCl (2.922g/L), and shaken for 7 days at 1000r.p.m. in a 37°C theromixer. The PFFs were subsequently aliquoted into 5ul tubes and stored in -80°C until used for in vitro assays.

### UV-visible spectroscopy of alpha-synuclein PFFs

Absorbance of alpha-synuclein PFFs in phosphate buffered saline (PBS) was measured on a NanoDrop One/One^C^ UV-visible absorbance spectrophotometer in the range 200-300nm. A dilution of 0.1μg/μL was prepared from a PFF stock solution of 5μg/μL, and added as a droplet (2μL) to the pedestal after different timepoints of ultrasonication (Branson CPXH Series Ultrasonic bath, 2.8L) (37, 40, 21°C). Data was collected with OneViewer Software.

### Atomic Force Microscopy (AFM)

AFM was performed with ScanAsyst Air tapping mode using an AFM Veeco, Multimode V. Samples were applied on mica and spread out to dry. Results were analysed with NanoScope Analysis 1.5 software.

### Microelectrode array (MEA) based electrophysiology

The spontaneous electrophysiological activity of the neural networks was recorded using an MEA2100 *in vitro* system together with the MEA suite software (Multi Channel System). The engineered neural networks were maintained on 60-electrode planar microelectrode arrays (MEAs) (60MEA200/30iR-Ti; Multi Channel Systems) with ring covers. Prior to seeding, the MEAs were briefly washed with 65% ethanol, incubated in sterile water and UV-treated. Subsequently, they were treated with foetal bovine serum for 30-60 minutes to make the surface hydrophilic, before being coated with 0.01% poly-L-ornithine (PLO) solution and L15/laminin. Each MEA (n=8) was seeded with 100,000 iPSC-derived neurons and kept in a standard humidified air incubator (5% CO^2^, 20%O^2^, 37°C). 50% of the media was changed every 3-4 days. Following 34 days of maturation, the PFFs were added to the neural networks. A 5μg/μL aliquot was thawed in room temperature and diluted in 245ul sterile PBS (0.1μg/μL). A water bath ultrasonicator (Branson CPXH Series Ultrasonic bath, 2.8L) was used to sonicate the PFFs for 1 hour (37, 40, 21°C), before 10uL of the PFF seeds (0.1ug/uL), or equivalent amounts of alpha-synuclein monomers or PBS, were added directly to the culture media. The MEA cultures were randomly assigned to the different test conditions: PFF group (n=4), PBS (n=2) and alpha-synuclein monomers (n=2). Network activity was sampled throughout the experimental period (7-minute recordings), where 5 baseline recordings, and 13 recordings after intervention was performed per MEA. To avoid inadvertent fluctuations in electrophysiological activity directly related to media changes, no recordings were performed in the first 48 hours following a media change.

### MEA data analysis

All data analysis, including the criticality analysis described in the following section, was carried out in MATLAB R2018b (The MathWorks, Inc.). The raw data from the MEA system was first bandpass filtered with a second-order Butterworth filter with a passband of 300 Hz to 3 kHz, and spike detection was performed on the filtered data using a threshold of 5 standard deviations below the median of the signal. After visual inspection of the filtered waveform-signal from each electrode, clear artefactual signals (outlier electrodes) were identified and removed from further analysis. A total of 11 such instances were identified, 9 of which were caused by the same electrode across multiple MEAs.

Four basic parameters were evaluated in an attempt to identify different functional behaviours in the different types of networks: the mean firing rate (MFR), inter-spike interval (ISI), population inter-spike interval (PISI), and cross-correlation (XC). All of the parameters were obtained from spike trains generated for each recording channel, where a spike train is given as a series of impulses with each impulse occurring at the time at which the peak voltage was recorded for each detected spike. The MFR for each recording channel was calculated as the total number of spikes detected on that channel divided by the total recording time. The MFR for a given network at a given time point was then taken as the average over all recording channels. The ISI for each recording channel was calculated as the average time interval between consecutive spikes detected on the same channel, and this was then also averaged over all channels for a given network at a given time point, excluding any intervals greater than 100 ms. The PISI was calculated by obtaining a population vector of the unique spike timings on all recording channels and averaging the intervals between them, excluding any intervals greater than 100 ms. This upper bound was selected based on previous reports of timing between successive spikes network-wide to provide an indication of information transmission in the network (40). The XC was obtained by computing the maximum autocorrelation-normalized magnitude of the cross-correlation of pairs of spike histograms for each pair of recording channels and averaging over all possible pairs. Spike histograms were obtained by temporally binning the spike trains with a bin size of 10 ms. The maximum lag considered in the XC calculation was 50 ms.

### Computational analysis of criticality

A flowchart showing the main steps of the criticality analysis can be found in Fig. 2. Preliminary analysis using the same method has been previously reported for one of the control networks from this dataset (63). Filtering and spike detection were first performed as described in the previous section (step 1, Fig. 2).

**Fig.2.**
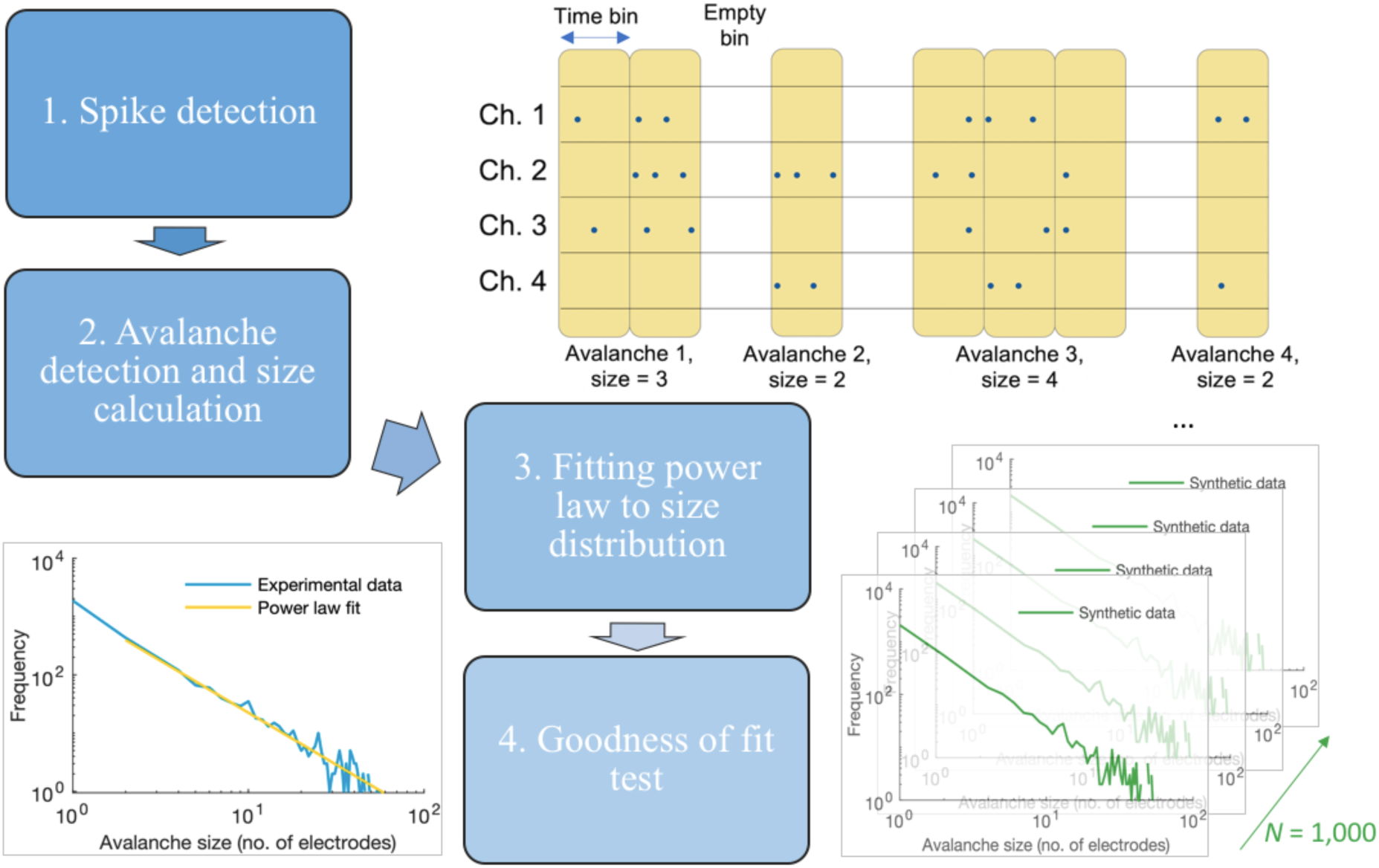
Step-by-step Criticality assessment.

Avalanche detection was then performed using the following procedure, based on the method originally described by Beggs and Plenz (45) (step 2, Fig. 2). The spike data were binned with a bin width of 1 ms, and avalanches were detected as any number of consecutive active time bins (bins containing one or more spikes) bounded before and after by empty time bins (bins containing no spikes). The avalanche size was computed as the number of active recording channels in the avalanche. (See the schematic in step 2 of Fig. 2 for an example.) The size probability distribution was then obtained by creating a histogram of the number of avalanches of each possible size (1 to 60 electrodes) and normalizing it with respect to the total number of avalanches.

As described by Beggs and Plenz (35), a hallmark of criticality is the avalanche size distribution following a power law. Thus, to determine whether or not the networks were in the critical state at a given time point, power law fitting was performed on the avalanche size distributions using the method described by (64). The fitted power law takes the form

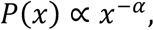

where *x* is the avalanche size, *P*(*x*) is the probability of an avalanche having size *x*, and *α* is the exponent of the power law. The fitting was performed for avalanche sizes ranging from 2 to 59 electrodes, following previous studies (e.g., (58)). Beggs and Plenz (45) originally reported *α* as taking a value of 1.5 in slice cultures, and this has been supported by other studies on dissociated cultures (e.g., (58)). When the fitted power law is plotted in log-log space, it appears as a line with a slope of -*α*. The goodness of fit was determined by generating *N* = 1,000 synthetic datasets from the fitted power law and computing the Kolmogorov–Smirnov (KS) distances for the empirical distribution and each of the synthetic distributions, where a greater KS distance indicates a poorer fit. The fraction *p* of synthetic distributions that had a KS distance greater than that of the empirical distribution (i.e., the fraction of cases where the empirical data were better described by the power law fit than were the synthetic data) was then calculated, and the fitting was rejected if *p* < 0.1, as suggested by Clauset et al. (53) as a more conservative threshold for the goodness-of-fit test. Thus, in the case where the fitting satisfied *p* ≥ 0.1, the network was presumed to be in a critical state.

### Immunocytochemistry

The engineered neural networks were fixed at room temperature with either 4% paraformaldehyde for 15 minutes, or 4% paraformaldehyde/4% sucrose/1% TritonX-100 (Sigma-Aldrich), as described in (62, 65) for protein extraction, at ranges between 10-20 minutes, followed by 3×15min washings with DPBS. TritonX-100 extraction should leave only insoluble inclusions, not showing any of the remaining presynaptic alpha-synuclein that has not converted to aggregates (15,20). Blocking was performed with a solution of 5% normal goat serum and 0.6% TritonX-100 in DPBS for 1 hour on a rotator at room temperature. Primary antibodies were subsequently applied overnight at 4°C, on a rotator, in a solution containing 2.5% normal goat serum and 0.3% TritonX-100. The following primary antibodies were used: rabbit polyclonal anti-alpha synuclein antibody 1:200 (ab131508, Abcam), mouse monoclonal anti-tyrosine hydroxylase antibody 1:300 (MA1-24654, Invitrogen), rabbit monoclonal anti-alpha synuclein (phospho S129) antibody 1:750 (ab51253, Abcam), chicken polyclonal anti-neurofilament heavy polypeptide 1:150 (ab46800, Abcam), and mouse monoclonal anti-beta-3 tubulin antibody 1:800 (ab119100, Abscam). The samples were then washed 3×15 min in DPBS at room temperature before being incubated in secondary solution containing 2.5% normal goat serum, 0.3% TritonX-100 and fluorophore-conjugated secondary antibodies 1:1000 (AlexaFluor 488, 568, 647, Life Technologies) for 3 hours. During the final 5 minutes of incubation, Hoechst was added at a final concentration of 1:10000. The samples were then washed 3×15 min in DPBS on a rotator. Some samples were also incubated with Phalloidin-iFluor 647 reagent – cytopainter 1:100 (ab176759, Abcam) for 20 minutes, before being washed 3×15min in DPBS again. Subsequently the samples were briefly washed in MilliQ-water, and mounted on Menzel glass-slides (Thermo Scientific) using FluorSave reagent (EMD Millipore USA).

### Transmission Electron Microscopy (TEM)

The neuronal cultures from two MEAs (one from the PFF group, one from the monomer control group) were detached from the surface of the MEAs by light suction using a 1000ul pipette and washed in DPBS, and immersed directly in 2.5% glutataldehyde without dissociation. The samples were subsequently stored at 4°C until further processing. In preparation for TEM, samples were gelatine embedded, dehydrated, infiltrated and blocked. A detailed description of the process can be found in the supplementary section.

Following processing for TEM, the embedded samples were sectioned (Ultramicrotome, Leica EM UC7) into 45-55nm thin sections, placed on grids, viewed with a Transmission Electron Microscope FEI Tecnai 12, and imaged with a Morada digital camera. Image processing was done using iTEM and Fiji.

## Results

### Formation and characterization of engineered neural networks on MEAs

After concluding the reprogramming protocol for human iPSC-derived neurons, the cells were seeded on MEAs and ibidi chips, where they spontaneously formed interconnections and extensive neural networks throughout the maturation period (**Fig.3**). Immunocytochemistry revealed neurons positive for beta-III tubulin, neurofilament heavy, and tyrosine hydroxylase in the engineered neural networks after 30 days of maturation. Importantly, the neural networks also expressed endogenous alpha-synuclein, which is a prerequisite for the induction of alpha-synuclein aggregation and pathology (**Supplementary Fig.S1**) (41).

**Fig.3.**
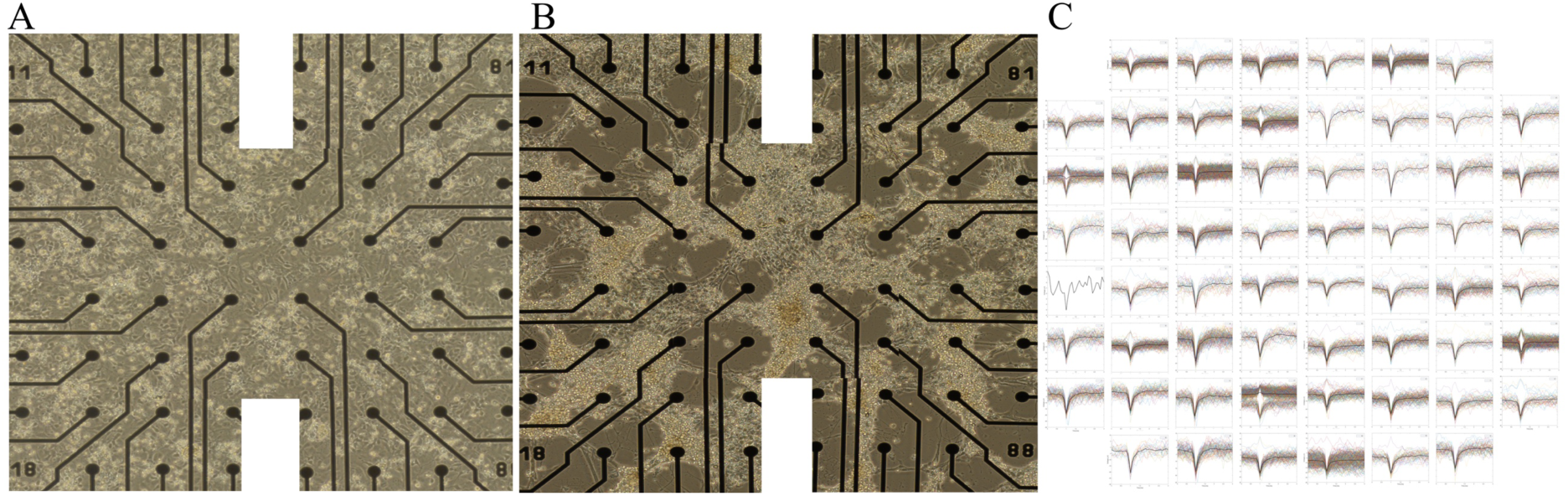
Formation and maturation of human neural networks on MEAs. **A**) shows a tiled microscopy image overviewing the electrode area of a newly seeded MEA (day 2 post seeding). **B**) shows how an extensive interconnected neural network has developed on the MEA surface 20 days post seeding. **C**) shows all of the individual spike shape cut-outs in colour (relative axis) obtained from each of the electrodes during a single recording session, demonstrating the electrophysiological activity obtained from a neural network as shown in **B**). The stronger black line within the spike shape cut-outs indicates the average spike shape for that electrode.

### Induction of alpha-synuclein pathology in neural networks

UV-visible absorbance spectra and AFM verified the breaking up of alpha-synuclein PFFs into smaller seeds by water bath ultrasonication, as a clear difference in both absorbance and structure of the PFFs was visible before and after sonication (**Supplementary Fig.S2**). Two weeks after the addition of sonicated PFF seeds, neural networks on ibidi chips were fixed and stained with the antibody for alpha-synuclein phosphorylated at S129 (ab51253) to visualize intracellular alpha-synuclein aggregates by immunofluorescence (**Supplementary Fig.S3**). Although consistent positive intracellular labelling by the S129 antibody was observed in the PFF treated neural networks, both perinuclearly and at distal neuronal sites, background staining and unspecific labelling was also consistently observed in the control conditions, even following TritonX-100 protein extraction, rendering the immunocytochemistry inconclusive.

### Verification of induced pathology in super resolution

Ultrastructural analysis of the neural networks collected from the MEAs showed evidence of perinuclear fibrillization in the samples from the PFF condition (**Fig.4A-D)**, but not in the samples from the monomer control condition (data not shown). Several fibrillous structures were also observed in the cytosol and within neurites of samples taken from the PFF condition (Fig.4**E-H**). Furthermore, an abundance of membrane-enveloped “inclusion bodies” in line with recent publications (66), were observed in the PFF condition, but not in samples from the monomer control condition (**Supplementary S4).**

**Fig. 4.**
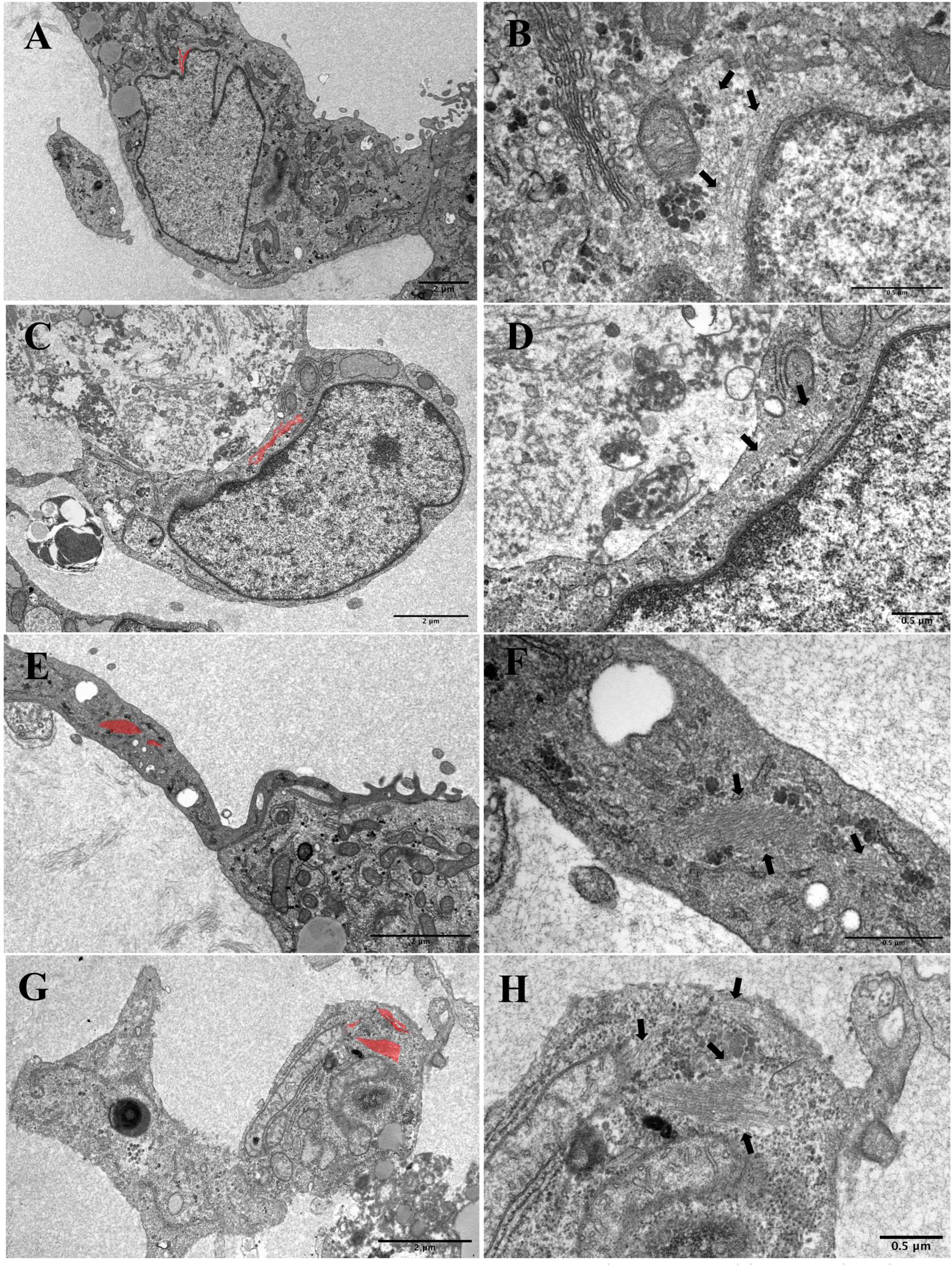
Fibrillization in PFF condition samples. **A-D)** Ultrastructural images showing perinuclear fibrillization in neural network samples from the PFF condition. (**A, C**) overview of single cell with intracellular features of interest highlighted in red (2μm scale bar). **B, D**) ultrastructure of perinuclear fibrils highlighted in panels **A, C** and indicated by black arrows (0.5μm scale bar). (**E**) overview image from the PFF condition showing fibrillization (highlighted in red) within a neurite, and **G**) fibrillization within the cytosol (2μm scale bar). **F, H**) shows the ultrastructure of the fibrils highlighted in panels E, G and indicated by arrows (0.5μm scale bar).

Furthermore, the ultrastructural analysis revealed a significant difference in observed necrotic and apoptotic elements in the extracellular environment surrounding neurons in the samples taken from the PFF treated condition and in the samples taken from the monomer control condition (t_14_=2,481, p<.05). In addition, the intracellular environment of single neurons revealed prominent autophagosomal and lysosomal vacuolization in samples from both the PFF treated condition and from the monomer control condition (with no significant differences between the conditions (t_15.549_=-.111, p>.05). (**Supplementary Fig.S5**). Representative overview images of samples used for ultrastructural analysis of extracellular necrotic/apoptotic elements, as well as intracellular autophagosomal and lysosomal vacuolization, can be found in the supplementary **(Fig.S6**,**S7).**

### MEA recordings and analysis

After 3 weeks of maturation on the MEAs, the baseline activity of the engineered neural networks (n=8) was recorded for 5 sessions until the point of PFF/monomer/PBS addition. After this point, 13 recordings (spanning across a total of 3 weeks) were made from the neural networks on each MEA (**Fig.5A**). Raster plots showing the spiking activity recorded from each individual MEA network during two common time points (recording 4 during the baseline period and recording 12 during the post PFF/monomer/PBS period) are displayed in **Fig.5BC**, illustrating both the developing activity patterns as well as the individual variability observed across MEAs.

**Fig.5.**
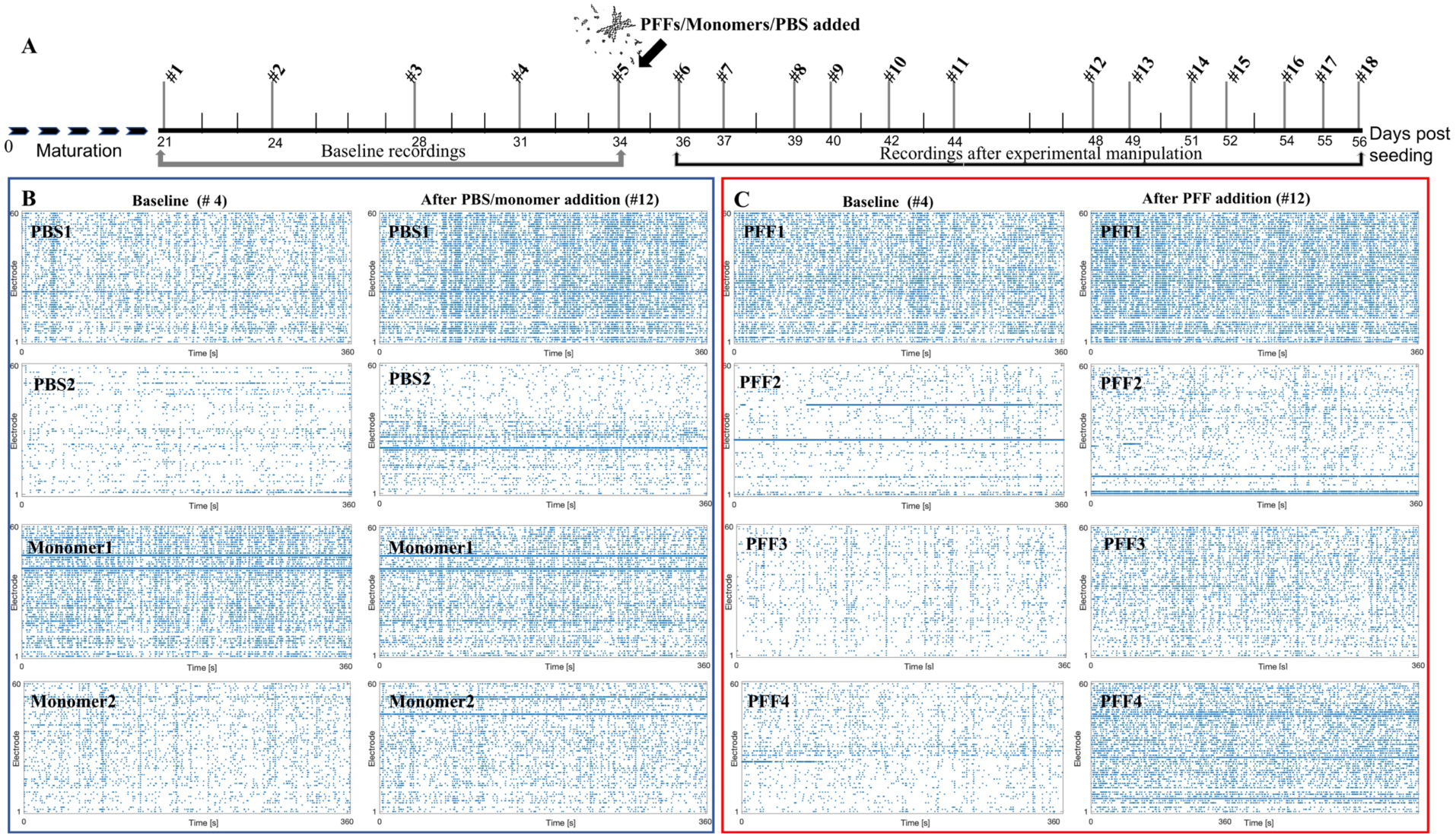
Microelectrode arrays (MEA) recording timeline and example raster plots from each neural network. **A**) depicts the MEA recording timeline, with # numbers indicating each recording time point and the axis numbers indicating the corresponding culture age (days post seeding). The arrow indicates the point of experimental intervention, where either PFFs, monomers or PBS were added to the neural networks after the 5^th^ baseline recording. Panels **B** and **C** show raster plots of the electrophysiological spiking activity obtained from each individual MEA neural network, with each blue dot representing a spike recorded at the indicated electrode. **B**) shows the spiking activity recorded from each of the control neural networks, two from the PBS condition (uppermost panels) and two from the monomer condition (bottommost panels), at a baseline time point (#4) (left), and at a post PBS/monomer time point (#12) (right). **C**) shows the spiking activity recorded from the experimental condition neural networks (n = 4), at the same baseline timepoint (4#) (left), and post PFF addition time point (#12) (right), as for the control condition MEAs displayed in panel **B**.

Four basic parameters were evaluated to observe how the networks matured: the MFR, ISI, PISI, and XC. The MFR describes the overall amount of activity in the network, and the ISI gives an indication of burstiness or the degree to which spikes from the same neuron occur in close temporal proximity. The PISI reflects network-wide spiking intervals and thus is expected to give an indication of connectivity or synchrony. Similarly, the XC describes the similarity between the spike trains from two recording channels and thus also gives an indication of functional connectivity or synchrony within the network.

**Fig. 6A** shows the mean baseline values of these parameters obtained for each group, and **Fig. 6B-E** shows plots of the time evolution of the parameters as percentages of the baseline values. The error bars represent the standard deviations among each group. One outlier recording from a neural network in the PFF group (MEA 16, time point 15) was eliminated because it had a high level of noise and appeared to yield many false positives in the spike detection, producing a spurious peak in the MFR and XC values. As shown in the results in **Fig. 6**, no apparent difference was observed among the evolution of these parameters, and thus no strong conclusions could be drawn about the difference in behaviour among the three groups.

**Fig.6.**
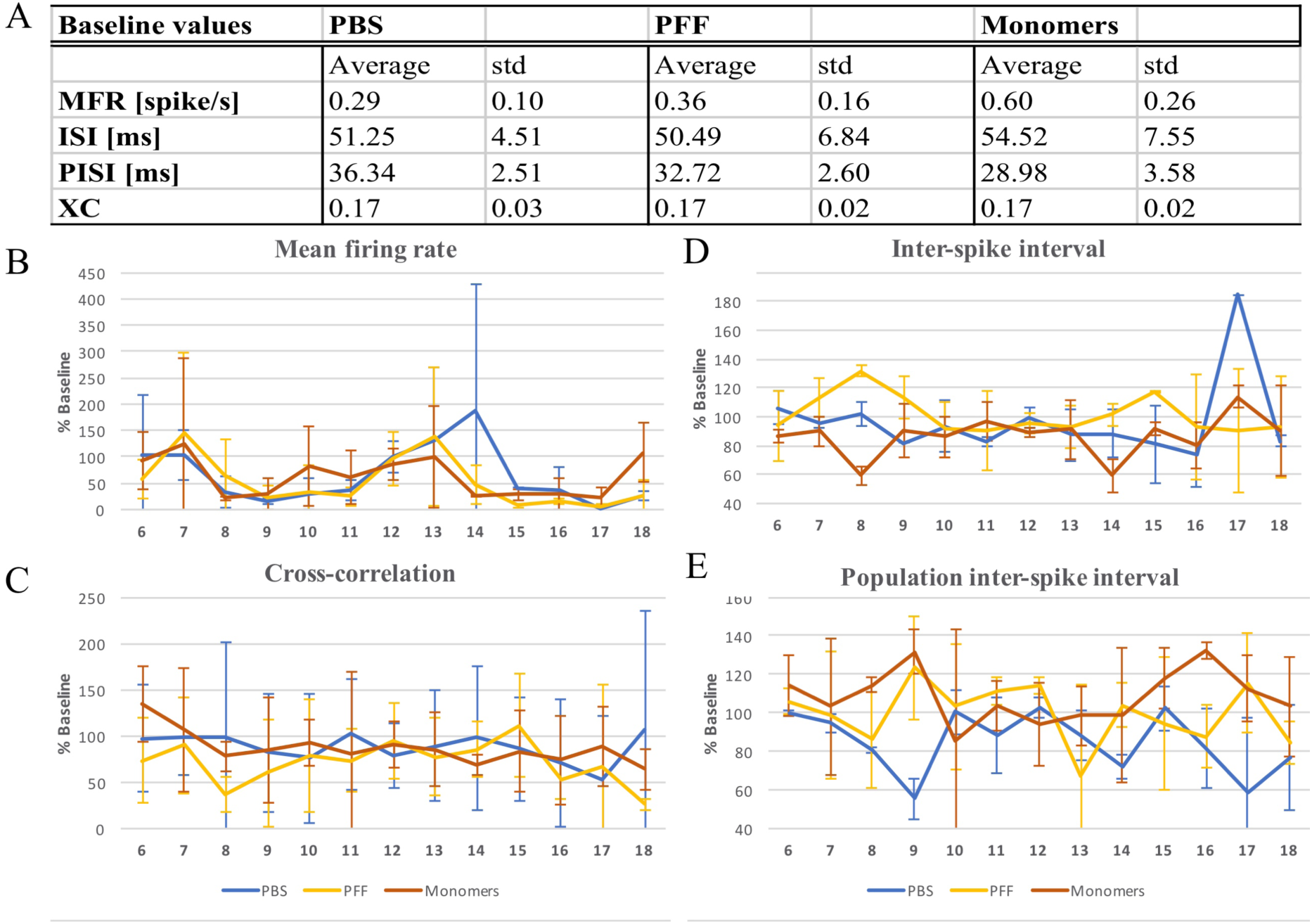
Descriptive electrophysiological values obtained throughout the recording period for all neural networks. **A)** table showing the average measures of the mean firing rate (MFR), inter-spike interval (ISI), population inter-spike interval (PISI), and cross-correlation (XC) measured across the 5 baseline time points for each group (PBS, PFF, and Monomer conditions), with standard deviations (std). **B-E**) shows a graphical representation of the MFR, XC, ISI, and PISI development, respectively, of all groups after the baseline period, with error bars representing the standard deviation across the networks in each group. Each value is given as a percentage of the baseline measures listed in table **A**).

### Assessment of Criticality

Criticality assessment of the 8 neural networks (2 monomer controls, 2 PBS controls, 4 PFF condition) revealed two clear outliers which were subsequently excluded, both of which were from the monomer control condition. One of these networks consistently displayed non-critical activity (during all 18 recording time points, both at baseline and following monomer addition), while the other network either displayed too few neuronal avalanches for criticality assessment or non-critical activity.

Recording time points where more than half of the neural networks did not exhibit enough neuronal avalanche activity for computational analysis of criticality have been omitted from the graphical representation in **Fig.7** (recording numbers 3, 8-11, 16-18). The criticality assessment at 4 baseline time points, as well as 6 time points following PFF addition are presented for 2 MEAs in the PBS condition (control), and 4 in the PFF condition (**Fig.7**). Analysis of criticality revealed fluctuating neural network states in both the PFF and PBS conditions. As can be seen from **Fig.7**, all neural networks (with the exception of network number 2 and 3 in the PFF condition) show probability size distributions of neuronal avalanche activity consistent with both critical and non-critical states during baseline measures, that is, before any perturbation. Furthermore, although some data points are missing (due to too few avalanches during the recording), most measurements during the baseline period are consistent with non-critical states (10/17 data points). However, after addition of alpha-synuclein PFF seeds to the neural networks in the PFF condition (represented by a black separation line in **Fig.7**), the majority of these perturbed neural networks (with the exception of PFF 4) mainly display critical activity states (11/17 data points). Contrary to this, the two neural networks in the PBS condition collectively display mostly non-critical activity states during these time points (6/9 data points). Together, these results suggest a difference in network criticality state between the groups after the point of perturbation, where the neural networks with PFF induced pathology largely display activity consistent with critical states, and the non-perturbed networks largely display activity consistent with non-critical states.

**Fig.7.**
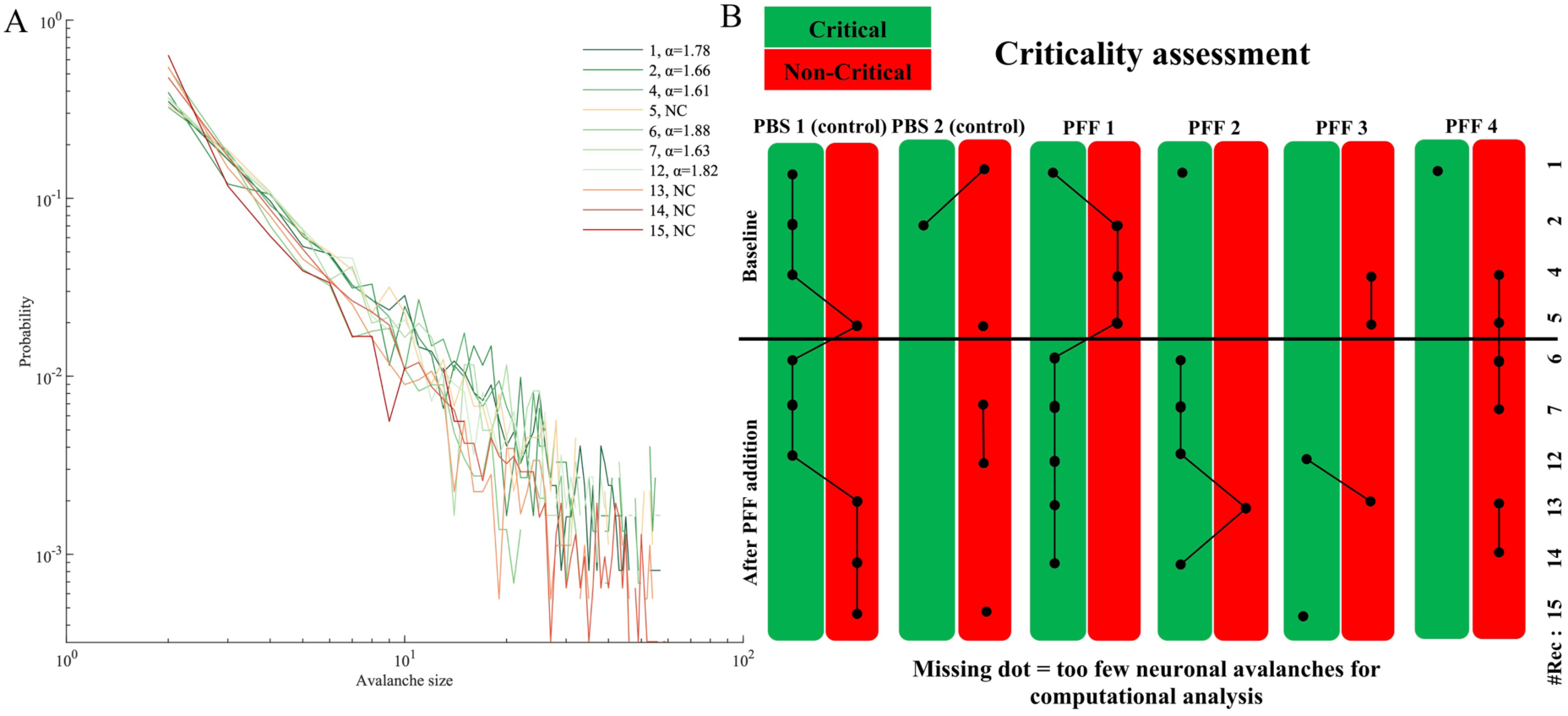
Criticality analysis. **A)** Shows the probablility distribution of the avalanche size for the 10 recording timepoints included (4 baseline, 6 after PBS addition) from the PBS 1 (control) neural network, with the power law exponent *α* values indicated for each time point where the power law fitting results indicated criticality. **B)** shows the cumulative criticality assessment for each of the 6 neural networks (2 from the PBS condition, 4 from the PFF condition) during the 10 recording time points included in the assessment. The point of perturbation (addition of PFF seeds, or PBS in the control condition) is indicated by the horizontal separation line. The green columns indicate neural network activity with avalanche size distributions following a power law distribution, consistent with a critical state (*p* ≥ 0.1), while red columns indicate a poor power law fit consistent with non-critical activity (*p* < 0.1). During the baseline time points, all neural networks (with the exception of PFF 2 and 3) fluctuated between critical and non-critical activity, with most data points being consistent with with non-critical states (10/17 data points). After the point of perturbation, the control neural networks (PBS 1 and 2) collectively display mainly non-critical activity states (6/9 data points), while the PFF neural networks mainly display critical activity states (with the exception of PFF 4) (11/17 data points), suggesting a difference between the groups in cirticality follwing pathology induction.

## Discussion

### Self-organized criticality

To investigate whether development of PD-related proteinopathy can be associated with network states of criticality, we induced alpha-synuclein proteinopathy in engineered human neural networks in vitro, and applied computational analysis to identify SoC in electrophysiological MEA recordings from the resulting network activity. Overall, our results point towards a difference in criticality state between the groups following perturbation, as the neural networks in the PFF condition largely ended up within the critical regime (10/17 data points), consistent with SoC, while the PBS controls largely ended up within the non-critical regime (6/9 data points) (**Fig.7**). SoC has been proposed as a mechanism that guides the spontaneous activity of developing neural networks into transient and homeostatically regulated patterns, or “meta-stable dynamics” (67, 68). These meta-stable dynamics are in turn part of the regular developmental trajectory of neural networks in vitro, and have been found to occur only in neural networks where the activity propagates within the critical mode (68). It is thus surprising that the networks with PFF induced pathology displayed neuronal avalanche activity largely consistent with SoC, while the control neural networks largely displayed non-critical activity. Nevertheless, these results could be indicative of a difference in developmental trajectory between the neural networks with PFF-induced pathology and the PBS control neural networks, highlight the potential relevance of SoC in unveiling functional alterations resulting from such an evolving pathology within the networks.

Evaluation of SoC through neuronal avalanche size distributions has been shown to provide a good representation of “damage spread” in perturbation experiments where identical replicas of the same system have different conditions and are investigated over time, even if the actual underlying dynamics are much more complicated (53). As can be seen from **Fig.5** summarizing the results of the standard electrophysiological analysis for the neural networks (MFR, XC, ISI and PISI) after perturbation, there is no clear trend separating the neural networks in the PFF condition from the networks in the control conditions, although the ultrastructural analysis revealed clear signs of induced pathology in the former (**Fig.4, S4-7**). This lack of a pathology expression in the functional activity of the perturbed neural networks is well in line with the aforementioned compensatory network mechanisms, such as homeostatic plasticity and circuit reconfiguration, which will work to maintain the functional capacity and present state of the network for as long as possible, effectively masking the developing PD-related pathology. However, as already noted, our results indicate a difference in network criticality state between the PFF and control group after perturbation, where the assessment of criticality should reflect the actual pathological development, whether it produces a linear, abrupt, or fluctuating change in the system dynamics, if enough time points and samples are incorporated (53).

Furthermore, the criticality analysis of neuronal avalanche activity revealed that the neural networks fluctuated between critical and non-critical states, both at baseline and after perturbation, for networks in both the PFF condition and the PBS control condition. Some other reports of in vitro neural network criticality (using dissociated rat cortical neurons) have found that most, but not all, of the neural networks investigated tend to stay within the critical regime after a certain point in their development (58, 68). The baseline activity of our human iPSC-derived neural networks mostly display activity consistent with a non-critical regime, suggesting a different developmental trajectory from networks derived from rodent primary neurons. This is indeed conceivable as the epigenetic and age-related imprint is removed through cell reprogramming through the iPSC stage (69, 70), resulting in a population of rejuvenated cells, from which our human neural networks were derived. Some studies indicate a slower development and maturation of neural networks derived from iPSCs compared to primary neurons (71-73), which could point towards a partial explanation of the largely non-critical activity observed at baseline in our neural networks. On the other hand, the non-homogenous population of cells represented within the iPSC-derived neural networks produces a more complex environment for development than pure neuronal cultures, which likely speeds up the developmental trajectory. For instance, the presence of astrocytes facilitates the formation and maturation of synaptic connections (74). However, as no other published study has investigated criticality in iPSC-derived neural networks, this remains to be elucidated, while other currently unknown influencing factors cannot be excluded at this point.

A more theoretical explanation for the large variability/ fluctuation observed in criticality state, both within and between the neural networks at baseline, can be found within the concept of “dirty criticality”. Dirty criticality, or “self-organized quasi criticality”, describes a mechanism which drags the activity back and forth around a stretched region of criticality, rather than being defined at a true point of criticality (which is needed to fully comply with standard SoC) (53, 75-78). This variant might thus contain more plausible models for biological neural networks, as neural network activity actually “hovers” around a region of criticality. Here, adaptive criticality (aSoC) models explicitly take into consideration the changing topology of the biological neural networks through dynamic parameters such as synaptic weight alteration, or link deletion and creation, thus encompassing structural network changes and local rewiring rules (75, 77-79). aSoC is thus based on a co-adaptive process between network architecture and dynamics (56, 80-82), meaning that the observed fluctuations in network criticality state during the baseline period could result from structural changes occurring in the networks (and *vice-versa*). Likewise, the fact that the neural networks in the PFF group largely ended up in a critical regime after pathology induction could be explained by the topological network changes caused by neuronal loss. Assuming that the networks were in a supercritical state prior to perturbation, a loss of connections would pull the network state towards criticality (56). Further corroborating this notion, a recent study by our group demonstrated that supercritical developing neural networks could be brought into a state of criticality through transient chemical disruption of the inhibitory-excitatory balance (83).

### Induction of pathology by alpha-synuclein PFF seeds

In a recent publication, Van den Berge et al., (13) showed that alpha-synuclein PFF seeds injected into the duodenum wall of a transgenic rat model induce an alpha-synuclein pathology which propagates transynaptically and bidirectionally through the parasympathetic and sympathetic nervous system to the brain stem in a pattern which recapitulates Braak’s hypothesis (6) of the development of a patterned pathological propagation in PD. This finding, together with the seminal demonstration of PFF induced pathology propagating from the gastrointestinal tract to the brains in rats (11), strengthens the growing suspicion of idiopathic PD originating from a yet unidentified pathogen capable of passing the mucosal barrier (84). Thus, our investigation of alpha-synuclein PFF induced pathology in human iPSC-derived neural networks is highly relevant and in line with the current direction of PD research.

As has been reported by several other studies investigating alpha-synuclein PFF induced pathology (85-90), we experienced issues with unequivocal identification of pathological alpha-synuclein aggregates using immunocytochemistry (**Fig.S3**), even after TritonX-1000 protein extraction, which should leave only insoluble inclusions (42, 62, 90). Although neural networks from the PFF condition consistently displayed positive immunolabeling of alpha-synuclein phosphorylated at S129, unspecific labelling and background staining were also observed in control conditions, rendering the immunoassays inconclusive. We therefore aimed to identify alpha-synuclein inclusions based on ultrastructural morphology and localization using TEM. In samples from the PFF condition, but not in samples from the monomer control condition, we observed several intracellular structures whose shape, size, and localization are consistent with alpha-synuclein aggregates found in previous studies (41, 85, 91)(**Fig.3, S4**). Furthermore, our neural network samples were analysed with respect to additional structural/morphological characteristics associated with reduced cell health and viability, and which are heavily linked to pathological intracellular aggregates (66). These include extracellular residual elements of apoptosis, necrosis and necroptosis (**Fig. S5, S6)**, as well as intracellular elements of autophagic and lysosomal activation **(Fig. S5, S7).** The latter is of particular interest as they are key regulators of cellular homeostasis, degrading and recycling proteins and cell constituents. As neurons are faced with disease related and aggregate-prone protein forms such as alpha-synuclein PFFs, this regulatory function becomes even more critical, as failure precipitates inclusion formation (92, 93). As pathological protein aggregation eventually saturates the autophagic machinery, the resulting imbalance in autophagic flux is believed to lead to neurodegeneration and cell death (94). This corresponds well with our results, as significantly more apoptotic and necrotic residues were found in the samples from PFF condition compared to the monomer control condition (**Fig.S5**). Furthermore, many of these apoptotic and necrotic elements showed signs of pathological fibril condensation (**Fig. S4**). As no significant difference in autophagic and lysosomal activation was found between the two conditions, these results suggest that most of the neurons affected by the PFF induced pathology had already succumbed to neurodegeneration and cell death at the point of ultrastructural analysis (38 days post perturbation).

## Conclusion

Our study shows that developing PD-related proteinopathy can be associated with network criticality states. Furthermore, the results suggest that induction of proteinopathy potentially changes the developmental trajectory of the neural networks in relation to SoC. Although the evolving pathology was not visible through common functional activity measures such as MFR, XC, ISI and PISI, a difference in overall criticality state suggests that there is a discernible difference between the PFF neural networks and the control neural networks after the point of perturbation, where the former largely displayed neuronal avalanche activity consistent with criticality, while the latter mainly displayed non-critical activity. To our knowledge, this first report associating network criticality state with induced proteinopathy in human iPSC-derived neural networks. This approach opens up exciting new avenues for identifying and understanding developing pathologies underlying neurodegenerative diseases.

## Abbreviations

PD: Parkinson’s disease
AD: Alzheimer’s disease
ALS: Amyotrophic lateral sclerosis
SNpc: Substantia Nigra pars compacta
MEA: multielectrode array
SoC: Self-organized criticality
PFF: alpha-synucelin pre-formed fibrils
iPSC: induced pluripotent stem cells
PBS: Phosphate buffered saline
AFM: Atomic force microscopy
TEM: Transmission electron microscopy
MFR: Mean firing rate
ISI: Inter-spike intervals
PISI: Population inter-spike interval
XC: Cross-correlation

## Acknowledgements

This work was supported by The Department of Neuromedicine and Movement Science, Faculty of Medicine and Health Sciences, NTNU; The Liaison Committee for Education, Research and Innovation in Central Norway (Samarbeidsorganet HMN, NTNU); and the SOCRATES project (NFR project agreement 270961). The TEM preparation was performed at the Cellular and Molecular Imaging Core Facility (CMIC, NTNU); The AFM and UV-vis spectroscopy were performed by Birgitte Hjelmeland McDonagh at the Norwegian Micro- and Nano-Fabrication Facility (NorFab), Norwegian University of Science and Technology (NTNU).

## Author contributions

VV designed the study, conducted the experiments, collected and analyzed the data, and wrote the paper; KH performed all electrophysiological data analysis, wrote parts of the paper, and contributed with critical discussion and interpretation of the findings; OR contributed with operationalizing the electrophysiological recordings and data analysis; WK created the PFFs, contributed with valuable feedback during the experimental process, and critically reviewed the paper; GB critically reviewed the paper; SN provided supervision and valuable discussion in relation to the electrophysiological data analysis and techniques; AS conceived and supervised the study, contributed to the study design, and critically reviewed the paper; IS conceived and supervised the study, contributed to the study design and interpretation of results, edited, and critically reviewed the paper.

## Competing interests

The authors declare no competing interests.

## Data availability

The datasets generated for this study are uploaded to Mendeley data repository and available in four separate folders as: Valderhaug, Vibeke D; Heiney, Kristine; Huse Ramstad, Ola; Nichele, Stefano, Sandvig, Axel; Sandvig, Ioanna (2020), “Crititcality as a measure of developing proteinopathy in biological human neural networks_Dataset 01-04”, Mendeley Data, V1, doi: 10.17632/x9w8×7xxr4.1 ; 10.17632/44d8wbxgr8.1 ; 10.1717632/zv3rktym53.1; 10.17632/m47bzbn425.1.

## Notes

### Competing Interest Statement

The authors have declared no competing interest.

