## Supplementary document for "Criticality as a measure of developing proteinopathy in engineered human neural networks"

Supplementary materials and results

### Materials and methods

*iPSCs to neural networks reprogramming protocol:*

| Day 0 - 4 | EB formation | | | |
|---|---|---|---|---|
| NIM - media | | Stock (mg/mL) | orking conc. | Total 100 mL |
| | DMEM/ F12 Media | 1X | 1X | 48ml |
| | Neurobasal | 1X | 1X | 48ml |
| | l-Glutamine | 200mM | 2mM | 1000ul |
| | NEAA (Non Essential Amino acids) | 100X | 1X | 1000ul |
| | 2-mercaptoethanol | 25mM | 12.5uM | 50ul |
| | Pen-Strep | 1X | 1X | 1000ul |
| | N2 supplement | 100X | 1X | 1000ul |
| **DAY 0 - 1** | | | | |
| | ROCK inhibitor Y-27632 | 10mM | 20uM | 200ul |
| | *SB43152 | 10mM (1000X) | 5uM | 50ul |
| | *LDN1931892 | 2mM (10 000X) | 100nM | 5ul |
| **DAY 1- 4** | | | | |
| | ROCK inhibitor Y-27632 | 10mM (50X) | 20uM | 200ul |
| | *SB43152 | 10mM (1000X) | 5uM | 50ul |
| | *LDN1931892 | 2mM (10 000X) | 100nM | 5ul |
| | * Shh (25II) | 10ug/ml (1000X) | 100ng/ml | 100ul |
| | *CHIR99021 (light sensitive) | 10mM | 0.7-0.8uM | 7.5ul |

| DAY 4-11 | Coat: Lam-111 | | | |
|---|---|---|---|---|
| NPM - media | | Stock (mg/mL) | orking conc. | Total 100 mL |
| | DMEM/F12 Media | 1X | 1X | 48ml |
| | Neurobasal Media | 1X | 1X | 48ml |
| | l-Glutamine | 200mM | 2mM | 1000ul |
| | NEAA (Non Essential Amino Acids) | 100X | 1X | 1000ul |
| | Pen-Strep | 1X | 1X | 1000ul |
| | **N2 supplement** | 100X | 0.5X | 500ul |
| **Day 4-9** | | | | |
| | **ROCK inhibitor Y-27632** | 10mM (50X) | 20uM | 200ul |
| | *SB43152 | 10mM (1000X) | 5uM | 50ul |
| | *LDN1931892 | 2mM (10 000X) | 100nM | 5ul |
| | * Shh (25II) | 10ug/ml (1000X) | 100ng/ml | 100ul |
| | *CHIR99021 (light sensitive) | 10mM | 0.7-0.8uM | 7.5ul |
| **Day 9 - 11** | | | | |
| | ***FGF8b** | 10ug/ml (1000X) | 50ng/ml | 50ul |
| | **B27 Supplement minus AO** | 50X | 1X | 2ml |

| DAY 11-16 | Dissociate using Accutase, replate on Lam-111 | | | |
|---|---|---|---|---|
| NDM - media | | Stock (mg/mL) | orking conc. | Total 100 mL |
| | Neurobasal | 1X | 1X | 97ml |
| | l-Glutamine | 200mM | 2mM | 1000ul |
| | NEAA | 100X | 1X | 1000ul |
| | Pen-Strep | 1X | 1X | 1000ul |
| Supplements | | | | |
| | **ROCK inhibitor Y-27632** | 10mM (50X) | 20uM | 200ul |
| | BDNF | 10ug/ml (1000X) | 20ng/ml | 200ul |
| | AA | 0.4mg/ml (1000X) | 0.4ug/ml | 100ul |
| | *FGF8b | 10ug/ml (1000X) | 50ng/ml | 50ul |
| | B27 Supplement minus AO | 50X | 1X | 2ml |

| Day 16------> Optional Maturation |  |  | PLO/lam |  |
|---|---|---|---|---|
| NDM - media | | Stock (mg/mL) | orking conc. | Total 100 mL |
| | Neurobasal | 1X | 1X | 97ml |
| | l-Glutamine | 200mM | 2mM | 1000ul |
| | Pen-Strep | 1X | 1X | 1000ul |
| | NEAA | 100X | 1X | 1000ul |
| | DAPT | 100mM | 10uM | 10ul |
| Supplements | B27 | 50X | 1X | 2ml |
| | Ascorbic acid (A) | 0.4mg/ml (1000X) | 0.4ug/ml | 100ul |
| | ROCK inhibitor (RI) | 10mM (50X) | 20uM | 200ul |
| | BDNF | 10ug/ml (1000X) | 20ng/ml | 200ul |

* Patterning factors

*TEM preparation of neural network samples:*

Gelatine embedding: All samples were washed for 2x10 min in 0.1M phosphate buffer, mixed 1:1 with gelatine (6% porcine, dissolved in 0,1M phosphate buffer) and incubated for 15-20min in 37-40°C. The samples were then centrifuged, cooled off, and post fixed with glutataldehyde. Surplus gelatine was cut away and the samples were cut and shaped into 1x1mm (or smaller) pieces.

Dehydration: The samples were washed 2x10min in 0.1M cacodylate buffer, and post-fixed in the dark for 1 hour in 2% $OsO_4$ + 1.5% potassium ferrocyanide in 0,1M cacodylate buffer. A series of washing and dehydration steps followed: 2x5min 0,1M cacodylate buffer, 2x5 min phosphate buffer, 1x10 min in 50% alcohol, 1x10 min in 70% alcohol, 1x10 min in 90% alcohol, 4x10 min in absolute alcohol, and 2x15min in ABS acetone.

Infiltration and embedding: 10ml epoxy was mixed with 0,15ml DMP-30. The samples were then placed in acetone and epoxy in the following dilutions: (2:1) for 2 hours, (1:1) for 2 hours, and (1:2) overnight. The samples were then placed in epoxy for 8 hours on a rotator, where the solution was replaced every 2 hours for better infiltration. The samples where then placed in plastic moulds, labelled, and polymerized at 60°C for 3 days.**)**

**Supplementary Result**

*Immunocytochemistr*

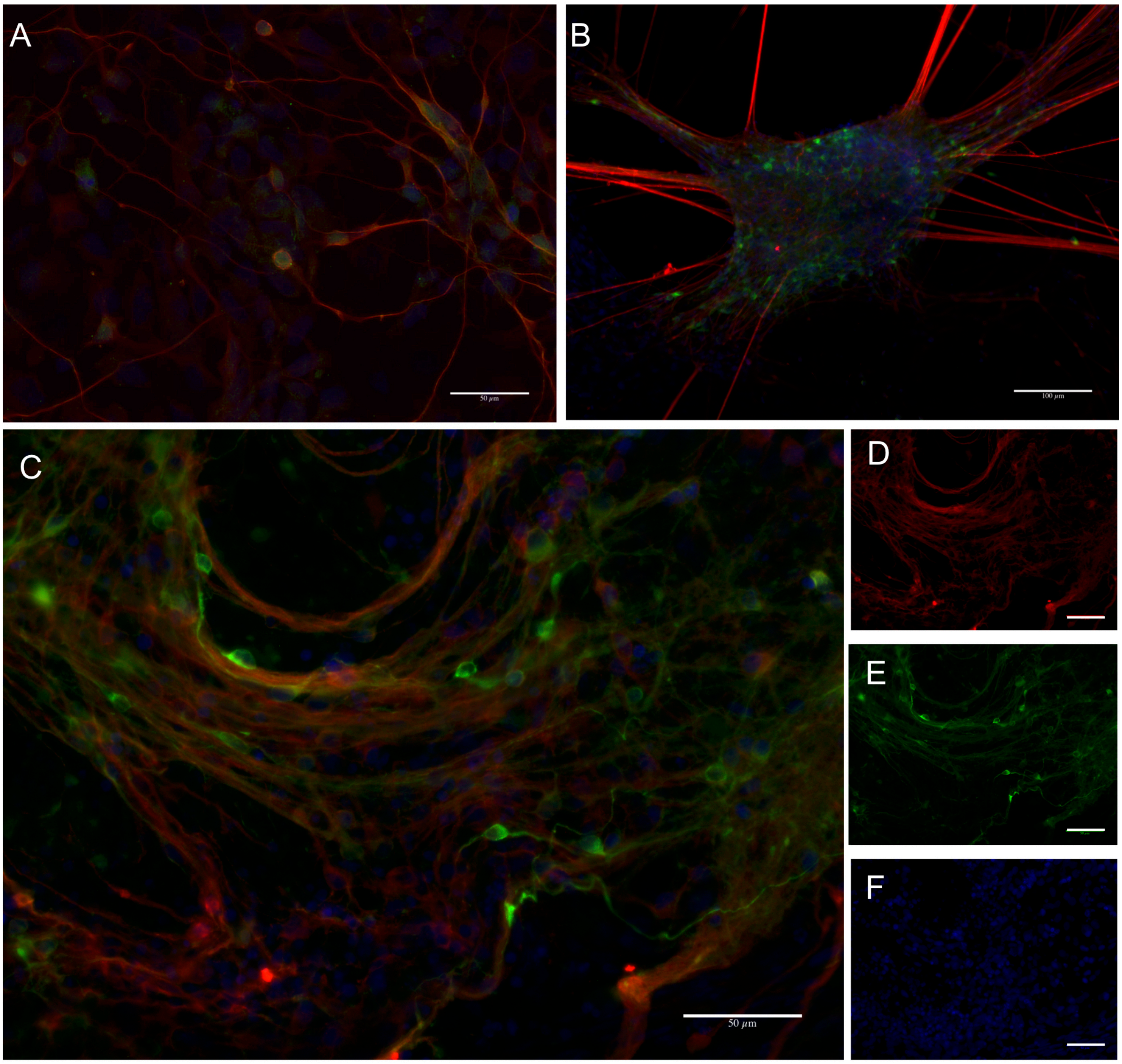

**Figure. S1 Antigenic profile of engineered neural networks after 30 days of maturation**. (**A**) Immunostaining with the neuron-specific cytoskeletal antibody beta-3 tubulin (red), the catecholamine neurotransmitter precursor antibody tyrosine hydroxylase (TH) (green), and counterstaining with hoechst (blue) (40X). (**B**) Immunostaining with the mature axonal antibody neurofilament heavy (red), TH (green) and counterstaining with hoechst (20X). (**C**). Image showing immunostaining with an antibody for endogenous alpha-synuclein (red) and TH (green), together with hoechst counterstain (blue) (40X). Images **D**, **E** and **F** are the single channel versions of **C**, showing endogenous alpha-synuclein, TH and Hoechst, respectively.

**Verification of alpha-synuclein PFF seeds**

UV-visible absorbance spectra and AFM of the PFFs showed a clear difference in both absorbance and structure of PFFs before and after water bath ultrasonication, indicating that the ultrasonication is effective in breaking the PFFs into smaller seeds.

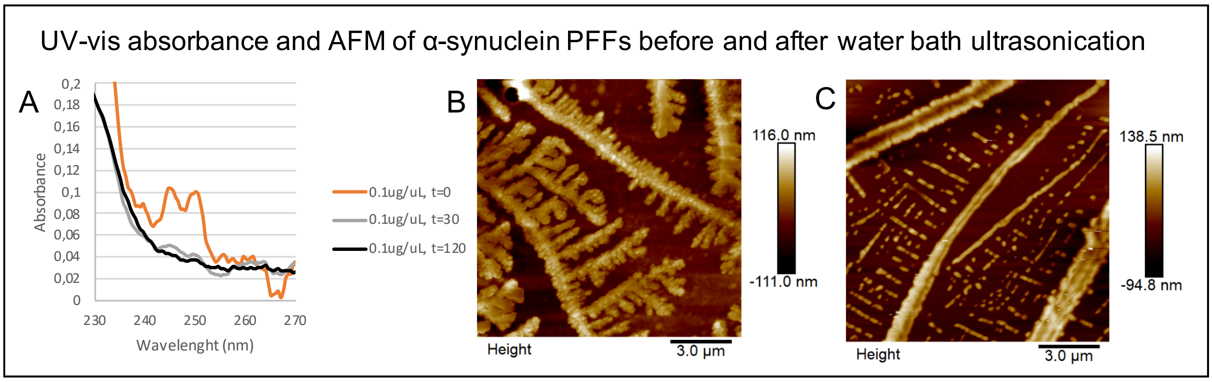


**Figure S2.**: **UV-visible absorbance spectra and atomic force microscopy (AFM) of pre-formed fibrils (PFFs). A**) UV-vis absorbance of α-synuclein PFFs show that before ultrasonication, the PFFs display a doublet absorption peak between $\lambda_{240}$ - $\lambda_{250}$. Absorption between $\lambda_{240}$ - $\lambda_{250}$ stems from $\pi \rightarrow \pi^*$ transitions of the pyrimidine and purine rings in nucleobases. After ultrasonication, the absorption at 240nm – 250 nm is reduced, which suggest less stacking of aromatic rings. AFM images **B**) before (t=0) and **C**) after (t=120 min) sonication show that treatment with ultrasonication disintegrates the branched structure of the PFFs.

**Verification of the presence of intracellular alpha-synuclein aggregates after PFF addition in engineered neural networks**

The antibody for alpha-synuclein phosphorylated at S129 (ab51253) was used to visualize intracellular aggregates by immunofluorescence in PFA-fixed and protein-extracted samples (TritonX-100) 2 weeks or more after addition of PFF seeds to the neuronal media. Although consistent positive intracellular labelling by the S129 antibody was observed in the PFF treated neural networks, both perinuclearly and at distal neuronal sites, background staining and unspecific labelling was also consistently observed in the control conditions (both in the DPBS and in the monomer control). Positive control labelling with the S129 antibody was observed at a range of different dilutions (1:100-1:750), with 3 different secondaries (488nm, 568nm, 647nm), for two different antibody batches, and after 10, 15, and 20 minutes' protein extraction by TritonX-100. Representative images of these results are shown in **S3**. The immunocytochemistry was thus inconclusive.

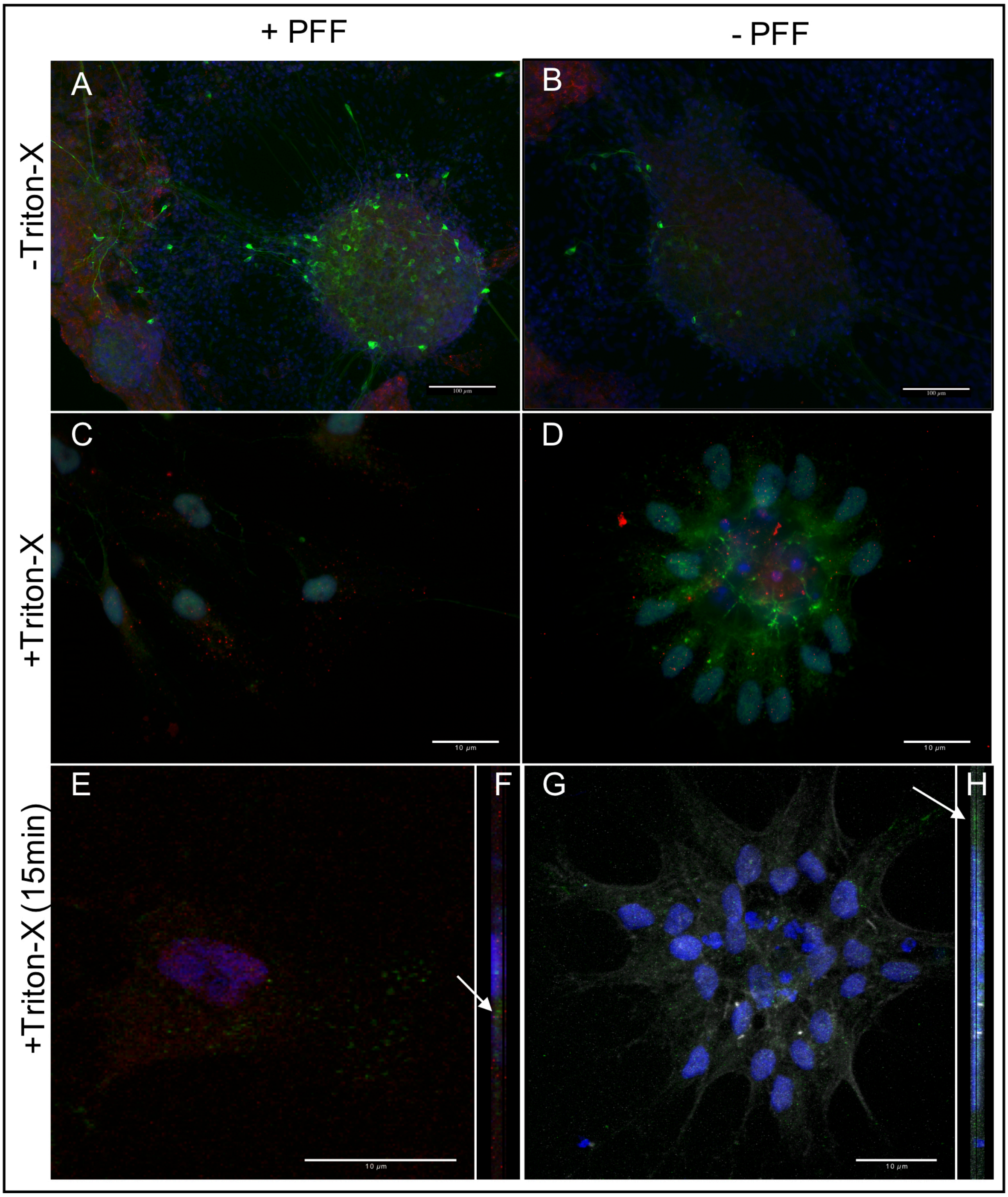


**Figure S3. Immunostaining of intracellular alpha-synuclein aggregates.** Immunofluorescent images displaying the detection of anti-alpha-synuclein phosphorylated at S129 antibody after different lengths of protein extraction in PFF cultures (**A,C,E** and **F**) and in control cultures (**B,D,G** and **H**). **A,B)** Immunocytochemistry of the mature engineered neural networks fixated without TritonX-100 extraction, displaying immunofluorescence with antibodies for tyrosine hydroxylase (green), for alpha synuclein phosphorylated at S129 (red) and with the nuclear counterstain hoechst (blue) (100um scale bar). **C,D)** Immunocytochemistry of cultures which were TritonX-100 extracted for 10 min prior to fixation, displaying immunofluorescent labelling with a phalloidin probe (green), the S129 antibody (red) and the nuclear counterstain hoechst (blue) (10um scale bar). **E and F)** Confocal image of a PFF treated culture which was TritonX-100 extracted for 15 min prior to fixation, displaying immunofluorescent labelling with a phalloidin probe (red), the S129 antibody (green) and the nuclear counterstain hoechst (blue) (10um scale bar) **F)** Side-view of **E)** created by confocal stacking (white arrow indicates intracellular localization of the S129 antibody). **G and H)** Confocal image of a control culture which was TritonX-100 extracted for 15 min prior to fixation, with immunofluorescent labelling with a phalloidin probe (grey), the S129 antibody (green) and the nuclear counterstain hoechst (blue) (10um scale bar) **H**) Side-view of **G**) created by confocal stacking (white arrow indicates intracellular localization of the S129 antibody).

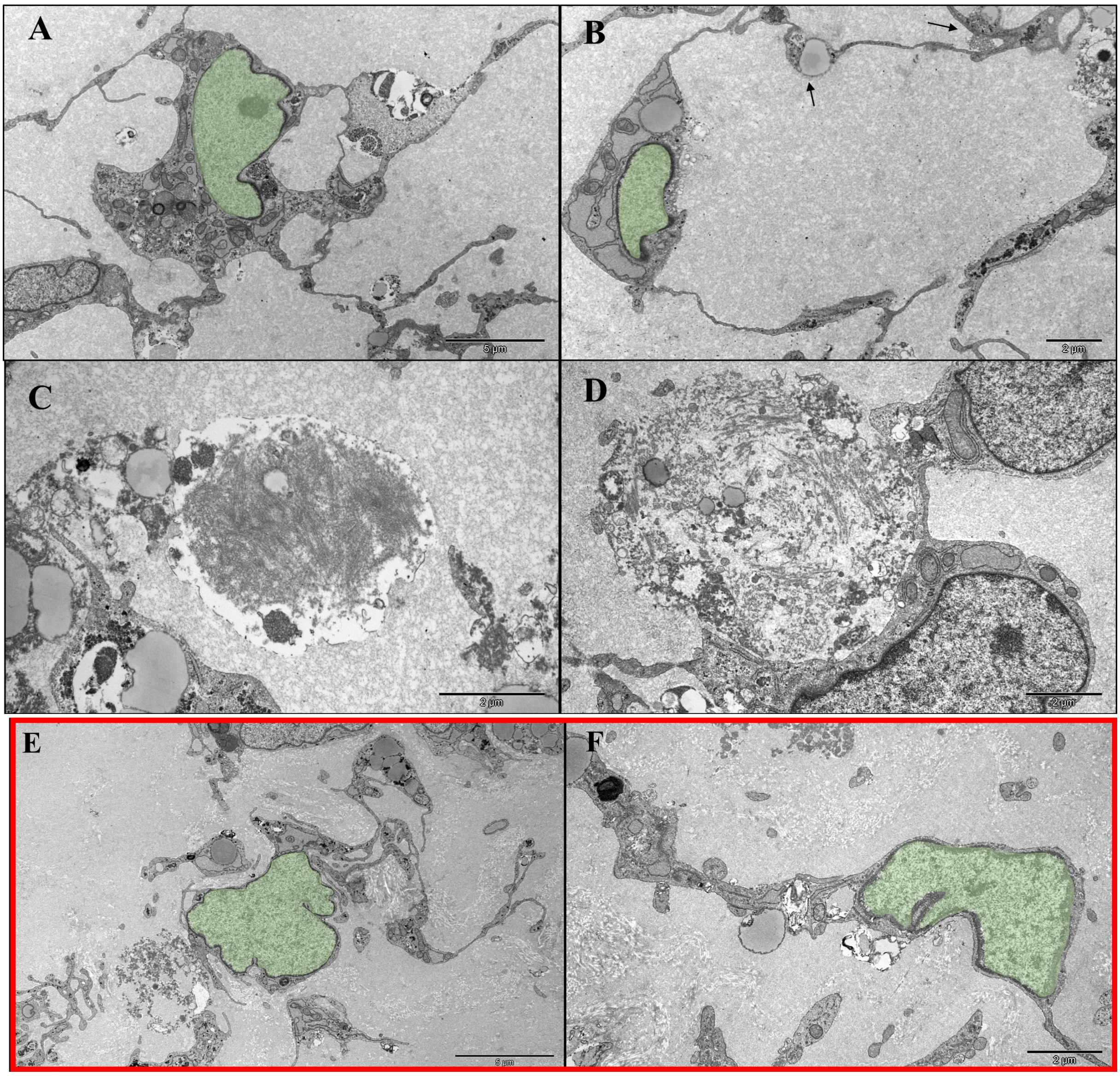


**Figure S4 Ultrastructural observations of fibrillization, inclusion bodies and neuritic atrophy A)** Ultrastructural sample showing an abundance of membrane-enveloped "inclusion bodies", both in the cytosol and within neurites of a neuron from the PFF condition (green = nucleus). **B**) shows a neuron from the PFF condition with signs of membrane degradation and atrophic neurites (arrow = swelling). **C**) and **D**) shows dead cells observed in the PFF condition with clear signs of fibril condensation. **E** and **F**) shows comparable neurons from the monomer control samples (green = nucleus).

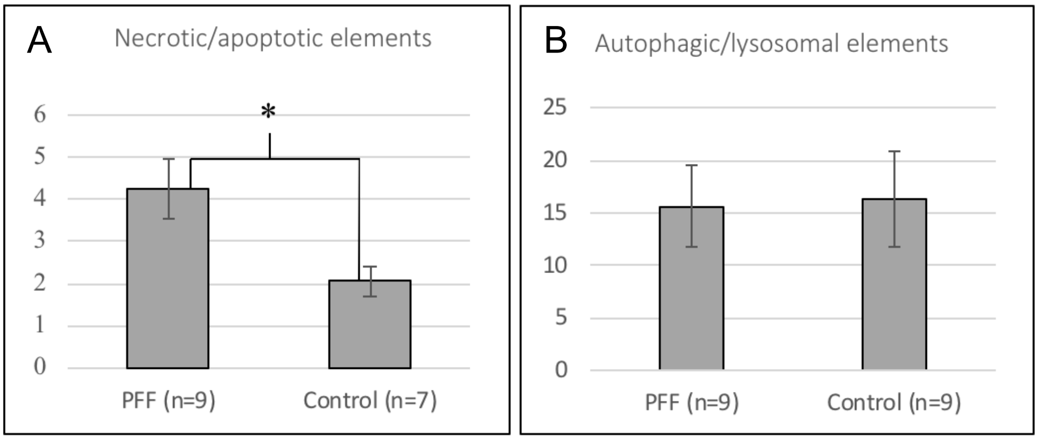


**Figure S5 Necrotic/apoptitic elements and autophagosomal/lysosomal activation.** Results from the ultrastructural image analysis, where the amount of necrosis-apoptosis (**A**) and autophagic/lysosomal elements (**B**) was compared between samples taken from the PFF treated condition and samples taken from the monomer control condition. The bar graphs display the results of the independent samples T-tests (group means with standard error of means). **A**) Mean number of apoptotic/necrotic elements surrounding single neurons in the PFF samples and the monomer control samples. n is the number of ultrastructural images analyzed for each condition, where the number of necrotic/apoptotic elements counted was divided by the number of morphologically intact neurons in the image (a total of 42 neurons from the PFF samples, and 39 from the monomer control samples). * significant at $p<.05$ **B**) No significant difference was found between the PFF samples and the monomer control samples in observed intracellular autophagic/lysosomal events. n denotes the number of single neurons analyzed.

**Fig.S6** shows representative overview images of samples used for ultrastructural analysis of extracellular necrotic/apoptotic elements. The regions of interest were selected based on the presence of a homogenous distribution of neurons.

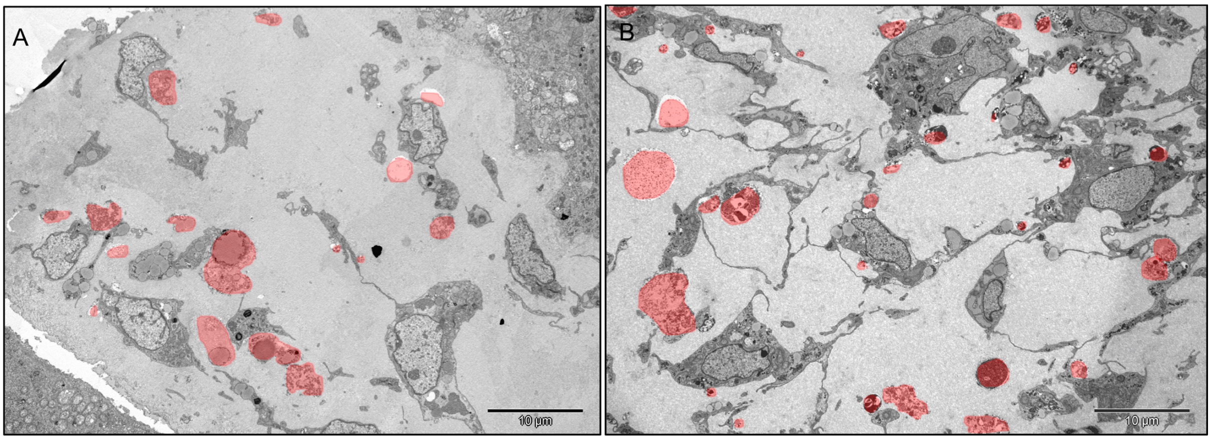


**Figure S6 Examples of ultrastructural images used for the assessment of extracellular necrotic/apoptotic elements:** A representative overview image from a monomer control sample (**A**) and a PFF treated sample (**B**), illustrating the observed differences in the extracellular environment surrounding the neurons. Red = highlighted residues of apoptotic/necrotic elements.

**Fig.S7** shows representative images used for the ultrastructural assessment of intracellular autophagic/lysosomal activation. The images analyzed were selected based on the presence of a number of neuron-specific ultrastructural features (axon/dendrites, synapses, nucleolus, shape and color of nucleus, as well as cytoplasmic electron density and heterochromatin condensation relative to other cell types identified in the preparation) (Peters and Folger; Rhodin 1974). Notice also the extracellular (filamentous) necrotic/apoptotic elements immediately adjacent to the neuron from the PFF condition displayed in **Fig.S5**.

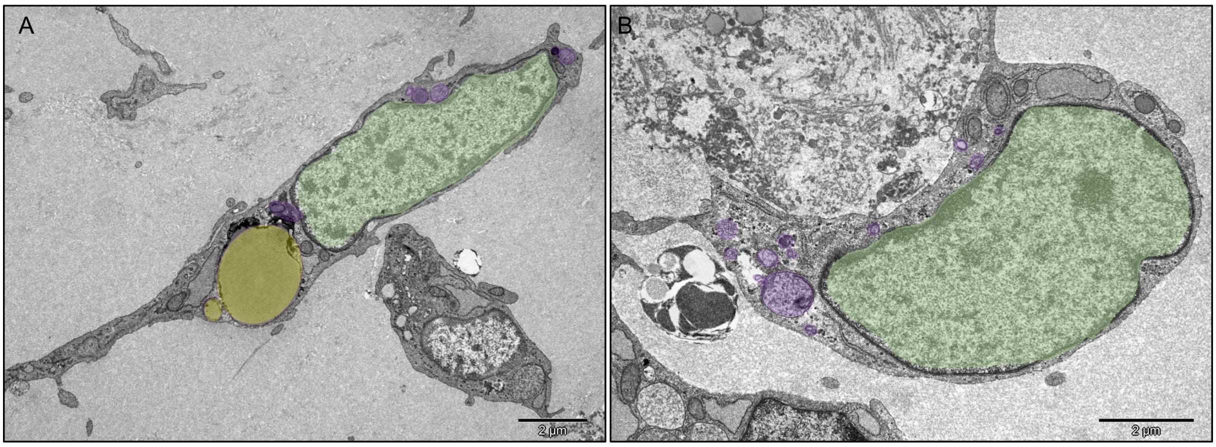


**Figure S7 Examples of ultrastructural images used for the assessment of autophagosomal/lysosomal activation:** Representative overview image of a single neuron from the monomer control samples (**A**) and from the PFF samples (**B**) illustrating the observed intracellular autophagic and lysosomal activity. Nucleus (green), purple autophagic/lysosomal activity (purple), lipid body (yellow).
